# Long-term mouse spinal cord organotypic slice culture as a platform for validating cell transplantation in spinal cord injury

**DOI:** 10.1101/2024.01.29.577615

**Authors:** Francesca Merighi, Sara De Vincentiis, Marco Onorati, Vittoria Raffa

**Affiliations:** Department of Biology, University of Pisa, Italy

## Abstract

Spinal cord injury (SCI) is an extremely invalidating condition with a severe physical and psychological impact. Resolutive cures are still lacking, due to its complex pathophysiology. One of the most promising regenerative approaches is based on stem cell transplantation to replace lost tissue and promote functional recovery. This approach should be explored better *in vitro* and *ex vivo* for safety and efficacy before proceeding with more expensive and time-consuming animal testing. In this work, we show the establishment of a long-term platform based on mouse spinal cord (SC) organotypic slices transplanted with human neural stem cells to test cellular replacement therapies for SCI. Standard SC organotypic cultures are maintained for up to 2 or 3 weeks *in vitro*. Here, we describe an optimized protocol for long-term maintenance for up to three months (90 days). The medium used for long-term culturing of SC slices was also optimized for transplanting neural stem cells into the organotypic model. Human SC-derived neuroepithelial stem (h-SC-NES) cells carrying a GFP reporter were transplanted into mouse SC-slices. 30 days after the transplant, cells still show GFP expression, and a low apoptotic rate, suggesting that the optimized environment sustained their survival and integration inside the tissue. This protocol represents a robust reference for efficiently testing cell replacement therapies in the SC tissue. This platform will allow researchers to perform an e*x vivo* pre-screening of different cell transplantation therapies, helping them to choose the most appropriate strategy before proceeding with *in vivo* experiments.

**SUMMARY:** In this paper, we provide a reproducible method to generate and maintain long–term spinal cord organotypic slices transplanted with neural stem cells as an *ex vivo* model for testing cellular replacement therapies.

## INTRODUCTION

Traumatic spinal cord injury (SCI) has devastating physical, psychological, and economic consequences for patients and caregivers. Globally, the incidence of SCI has increased over the last 30 years, with about 6.2 million total SCI patients in 2019 (*Ding, W. et al., 2022*). Many attempts have been made to promote axonal regeneration in SCI with different approaches (*Yang, B. et al., 2020*). Some beneficial effects were demonstrated by increasing the intrinsic growth potential of neurons through modulation of gene expression (*Liu, K. et al., 2010*) or administration of growth factors (*Anderson, M. A. et al., 2018*) and by the formation of neuronal relays between proximal and distal neurons in the injury site through cell replacement therapies. In particular, the interest in the use of cell therapies is growing exponentially (*De Freria, C. M. et al., 2021*), since transplanted cells can play many roles, including providing trophic support, immune modulation, regenerating lost neural circuits through induction of plasticity or cell replacement or axon remyelination (*Badner, A. et al., 2017*). Autologous or allogenic cells such as mesenchymal stem cells (NCT01676441, NCT03308565, NCT03521323; NCT03505034; NCT02481440), Schwann cells (NCT01739023, NCT02354625) and olfactory ensheathing cells (NCT01231893) were tested in SCI patients. Recently, the main effort in the field concentrated on human neural stem/progenitor cells (NSCs/NPCs), in particular those derived from induced pluripotent stem cells (iPSCs) (*Assinck, P. et al., 2017*). Studies published to date have shown that NSCs/NPCs transplantation for SCI is relatively safe (*Nógrádi, A. et al., 2011; Pereira, I.M. et al., 2019*). No significant increase in the rate of serious adverse events was reported, suggesting that the therapy is well tolerated (NCT02163876, NCT01321333, NCT 01772810) (Levi, A. D. *et al.*, 2018; Curtis, E. *et al.*, 2018). Studies suggest that NSCs/NPCs modulate the astrocyte response (*Ishii, K. et al, 2006*), promote the secretion of pro-regenerative factors (*Zhang, Y. W. et al., 2006; Faulkner, J. & Keirstead, H. S., 2005*), replace missing neuronal cells in SCI *(Kadoya, K. et al., 2016; Dell’Anno, M. et al., 2018)*, enhance oligodendrocyte and neuronal differentiation *(Kumamaru, H. et al., 2013)*, promote remodelling of neural circuitry and improve remyelination *(Duncan, I. D. et al., 2009)*. However, studies that support the differentiation of transplanted cells into functional neurons are still poor. Indeed, transplanted cell survival and differentiation in the injured spinal cord is low *(Wu, S., FitzGerald, K. T. & Giordano, J., 2018*), possibly because transplanted cells take several weeks, even months, to differentiate *in vivo*. Additionally, current studies have not elucidated many biochemical, molecular, cellular, and functional aspects of cell replacement therapies. In this context, simple, fast, and cost-effective models are required to study the mechanisms of cell engraftments, the ability of engrafted cells to proliferate, differentiate into specific types or subpopulations of cells, and form synapses with resident neurons. Integrating histological studies into electrophysiological recording and transcriptome and proteome profiling is necessary for a full comprehension of the molecular cascade occurring after cell transplantation. This certainly will speed up the design and validation of new replacement therapies in pre-clinical models and clinical studies. Indeed, up to now the use of rodents, large animals, and non-human primates has been worthwhile for elucidating many cellular processes after transplantation (*Nardone, R. et al., 2017*). However, due to the high cost, the high ethical impact, as well as the complexity of the organism, their use is often not straightforward or not totally adequate to unravel biochemical and molecular processes. In addition, they may present many disadvantages correlated with biological differences, both inter-species - *i.e.* metabolism -, and intra-species differences - *i.e.* sex, age - and associated with external factors like stressful situations, that could alter the outcome of an experiment and their predictability in terms of therapeutic translation to humans (*Hartung, T. et al., 2008; Shanks, N., Greek, R., Greek, J. et al., 2009; Dawson, T., Golde, T., Lagier-Tourenne, C. et al., 2018*). Thus, many groups employ 2D *in vitro* cell culture, and *ex vivo* organotypic slices (*ex vivo* cultures), in addition to animal models. 2D cell culture is the most commonly used system for studying specific biological processes at a single-cell and/or cell population level. Nevertheless, monolayer cell cultures don’t reflect the complexity found in a whole organism. The lack of tissue structures and the physiological environment do not allow the 2D culture system to completely emulate key structural, morphological, and functional aspects of the investigated tissue *(Hayden, P. et al., 2021; Jensen, C. et al., 2020; Mirbagheri, M. et al., 2019).* Organotypic cultures can overcome some of these issues. Organotypic models are based on explanting a fragment of a tissue or organ and maintaining it *ex vivo* for a limited period (*Gahwiler, B. et al., 1981*; *Stoppini, L., Buchs P., Miller, D., et al., 1991*). The slices of the explanted tissue have to be generated with a precise thickness that allows nutrients to reach easily almost all the cells in the slices. They can be generated from various regions of the central nervous system, like the hippocampus, hypothalamus, cerebellum, thalamus, cortex, substantia nigra and striatum, and spinal cord (*Pandamooz, S. et al., 2016*). Organotypic cultures present the advantage of retaining the tissue architecture, the spatial distribution of cells, the cellular diversity, and the environment (*i.e.* extracellular matrix composition) of the organ of origin. Moreover, they preserve the original neural activity, connections between cells, and in particular short-distance circuits after the explant. These aspects provide some advantages for *ex vivo* cultures with respect to both monolayer cultures and animal models. They still retain key tissue features found *in vivo* but with the reduction of the costs and the possibility to perform different types of molecular, cellular, and functional experiments with an accurate regulation of the culture environment’s parameters (*Fuller L. and Dailey M.E. et al., 2007; Gertz C.C. et al., 2014; Ballerini, L. and Galante, M., 1998; Avossa, D. et al., 2003; Lossi et al., 2018*; *Nogueira, G.O. et al., 2022*). Organotypic slices can also be exploited to develop models for different neurological disorders by resembling key histopathological features of specific conditions (*Qi, X. et al.; 2019*). Moreover, the retention of the original multicellular and tissue environment renders them appropriate platforms for drug screening and for testing neuroprotective and neuro-regenerative molecules and materials. In this work, we propose the use of SC organotypic cultures as a model to optimize NSCs transplants. This is not trivial, since optimal culturing conditions are required to guarantee the survival of both the host (SC tissue) and the transplant (NSCs) for weeks. Different research groups engrafted in organotypic cultures - both brain-derived and SC-derived – various types of cells. Most of the works showed the transplantation of mesenchymal stem cells (*Park, H. et al., 2010*; *Charriere, K., Risold, P., Fellmann, D. et al., 2010*; *Jeong, D. et al., 2011*), olfactory ensheathing cells (*Riggio, C. et al., 2013*) or NSCs (*Kamei, N. et al., 2004; Kamei, N. et al., 2007; Hamasaki, T. et al., 2007; Kim, H. et al., 2010; Liu, X. et al., 2014; De Vincentiis, S. at al., 2023*) and evaluated the interactions of engrafted cells with the host cells, the survival of the whole system, and whether the transplanted cells differentiated into neurons or neuron-like cells inside the *ex vivo* tissue environment (*Abouelfetouh, A. et al., 2004; Charriere, K., et al., 2010; Jeong, D. et al., 2011*). Some of them evaluated also the regenerative potential after transplant, observing the axonal growth inside the tissue (*Abouelfetouh, A. et al., 2004; Hamasaki, T. et al., 2007; De Vincentiis, S. et al., 2023*), the myelinating ability of engrafted precursors of oligodendrocytes (*Sypecka, J. et al., 2015*), the migration of engrafted cells into the host tissue (*Tanvig, M. et al., 2009*), and whether transplanted cells released factors pushing towards a pro-regenerative environment (*Park et al., 2010)*. One limitation of the current studies is that they do not explore the engraftment for a long-term period. Considering that NSCs seem to require several weeks to differentiate *in vivo* (*Tennstaedt, A. et al., 2015; Vogel, S. et al., 2019*), this study focuses on how to generate and maintain long-term mouse SC slices for up to 90 days. Slices were found to retain their original anatomical structure and to maintain a low and stable apoptotic rate over time and high cell viability. We observed diffuse expression of neuronal markers RNA binding fox-1 homolog 3 (RBFOX3) and neurofilament light chain (NFL), with the latter showing an increasing trend of axonal sprouting around the slices over time, attesting to their healthy condition. Moreover, we successfully transplanted into SC-slices, GFP-expressing human SC-derived neuroepithelial stem (h-SC-NES) cells at the first stages of neuronal differentiation. The neural stem cell graft into slices was maintained for 30 days after transplant and cells showed GFP expression for all the period in culture. The apoptotic rate of cells at day post-transplant (DPT) 30 was also found to be in line with respect to the apoptotic rate value observed at DPT 7 in the same cells (*De Vincentiis, S. et al., 2023*). Cells seemed to engraft into the tissue environment and survived up to several weeks.

In summary, our data demonstrate that it is possible to maintain in culture SC organotypic cultures for 3 months, without compromising their original cytoarchitecture and the tissue environment. Most importantly, they can be exploited to test NSCs-based cell therapies, before proceeding with an *in vivo* experiment, reducing the costs and the experimental time. In the next pages, we illustrate in detail all the passages to generate mouse SC organotypic slices and how to maintain them for long-term periods. Moreover, we explain deeply how to perform cell transplantation into the slices and how to maintain them for downstream molecular and cellular assays.

## PROTOCOL

Ethical Statement: Animal procedures were performed in strict compliance with protocols approved by Italian Ministry of Public Health and the local Ethical Committee of University of Pisa, in conformity with the Directive 2010/63/EU (project license no. 39E1C.N.5Q7 released on 30/10/2021). C57BL/6J mice were kept in a regulated environment (23 ± 1 °C, 50 ± 5% humidity) with a 12 h light–dark cycle with food and water *ad libitum*.

All NES cell works were performed according to NIH guidelines for the acquisition and distribution of human tissue for bio-medical research purposes and with approval by the Human Investigation Committees and Institutional Ethics Committees of each institute from which samples were obtained. Final approval from the Committee on Bioethics of the University of Pisa was obtained (Review No. 29/2020). De-identified human specimens were provided by the Joint MRC/Wellcome Trust grant (099175/Z/12/Z), Human Developmental Biology Resource (www.hdbr.org). Appropriate informed consent was obtained, and all available non-identifying information was recorded for each specimen. Tissue was handled in accordance with ethical guidelines and regulations for the research use of human brain tissue set forth by the NIH (http://bioethics.od.nih.gov/humantissue. html) and the WMA Declaration of Helsinki (http://www.wma.net/en/30publications/10policies/b3/index.html).

### 1. Preparation of solutions and equipment for SC isolation and culturing

#### 1.1 Coating of culture insert-membranes

1. Prepare the coating solution: a water solution with 0.1 mg ml^-1^ collagen, 0.01 mg ml^-1^ poly-L-lysine, and 0.01 mg ml^-1^ laminin.
2. Place each insert membrane into 35 mm dishes or a 6-well plate.
3. Add to the top of the membrane 1 ml of coating solution: incubate the solution for 4 h at room temperature (RT); then, remove it and let the membrane dry overnight (ON). Store the membrane at 4°C until their use.

NOTE: All the passages should be performed in sterile conditions. Membrane coating should be done no more than one week before their use.

#### 1.2 Media preparation: organotypic medium (OM), dissection medium, graft medium (GM)

1. *Organotypic medium (OM)* is composed of minimum essential medium (MEM) 25% of horse serum, Hanks’ Balanced Salt Solution (HBSS) and N-2-hydroxyethylpiperazine-N-2-ethane sulfonic acid (HEPES), 6.3 mg mL^−1^ of d-glucose, 1% penicillin, 1% streptomycin, and 0.1 μg mL^−1^ Glial cell-derived neurotrophic factor (GDNF).
2. *The dissection medium* with is composed of D-glucose 6.5 mg mL^−1^ in Dulbecco’s phosphate-buffered saline (DPBS).
3. *The graft medium (GM)* is composed of Neurobasal, N2 (1:100), B27 (1:50), Insulin (10 μg mL^−1^), Glutamax (1:100), 1% penicillin, 1% streptomycin, and 0.1 μg mL^−1^ GDNF (medium optimized from *Onorati, M. et al., 2016)*.

NOTE: solutions should be prepared in sterile conditions and just before their use (one day before or the same day of the experiment).

#### 1.3 Materials preparation for surgery

1. Prepare in the biosafety cabinet the surgical instruments: one large scissor, one micro-scissors, two straight tweezers, and two curved tweezers.
2. Prepare in the biosafety cabinet the chopper instrument by equipping it with a blade to cut the SC into slices. The blade should be placed precisely perpendicular to the cutting deck to perform the cutting correctly.
3. Prepare two plastic Pasteur pipettes, necessary to move isolated SC and the slices.

NOTE: It is necessary to sterilize all the instruments with ethanol 70% and UV (one cycle of 20 min) just before their use, to preserve culture sterility.

### 2. Isolation of mouse SC and slices generation

#### 2.1 Isolation of mouse SC organotypic slices

1. Decapitate P3 mouse pups with large scissors at the level of the foramen magnum.
2. Through a midline laparotomy with the micro-scissor, isolate the lumbar region of the backbone from the rest of the body and put it in cold dissection medium.
3. Using a dissection stereomicroscope, cut with the micro-scissor the backbone along the sagittal axis and using straight tweezers remove gently the SC from the backbone cavity.
4. Peel off carefully the meninges from the isolated lumbar region of the SC by using straight tweezers.
5. Incubate the isolated SC lumbar region into cold and fresh dissection medium for 10-15 min before proceeding with the subsequent step.

#### 2.2 Slice generation

1. By using one plastic Pasteur pipette, place the SC lumbar region on the cutting deck of a chopper instrument, perpendicularly to the blade. NOTE: Correct positioning of the SC with respect to the blade and of the blade with respect to the cutting deck is essential to properly generate SC slices.
2. Remove the residual dissection medium on the deck around the SC with the help of the Pasteur pipet and sterile absorbent paper. Set the cutting step of the chopper instrument at 350 µm and proceed with SC automated sectioning.
3. Collect the slices into a Petri dish with fresh dissection medium and incubate them for 15 min.
4. During slices incubation, wash three times the culturing insert-membranes with OM. Then, leave 1ml of OM at the bottom of each insert-membrane.
5. Seed the slices on the conditioned insert-membranes with the preferred orientation and in the desired position. Remove excess of medium to allow slices to better adhere to the membrane surface. NOTE: After the incubation in the dissection medium, slices are collected with a plastic Pasteur pipette and then put on the insert membrane. Seed on the membrane the desired number of slices. Then, remove the excess of dissection medium, and move and orient the slices with the straight tweezer. This operation should be performed gently to avoid tissue or insert damage.
6. After 30 min incubation at 37°C, change the medium: remove the spent medium and add 1ml of fresh one, supplemented with GDNF. Incubate the slices at 37°C. Refer to the first day in culture as day *ex vivo* (DEV) 0.

### 3. Long-term culturing of organotypic slices

1. Slices are maintained in culture at 37°C until desired time points.
2. Change the spent medium with fresh one at DEV 1 and then every other day, *i.e.* DEV 3, DEV 5…etc.
3. At DEV 3 switch the medium to the CCM to create the appropriate environment for transplanting stem cells the day after. NOTE: Add fresh GDNF every day until DEV 7 and then add it only every other day when the medium is changed.

### 4. Human neural stem cell culturing

#### 4.1 Cell maintenance, pre-differentiation and differentiation

h-SC –NES cells were maintained in culture in presence of growth factors (FGF-2 and EGF). Before the transplantation, a pre-differentiation protocol is performed by removing the growth factors for 7 days. The differentiation is supported by adding neurotrophin supplements (*e.g.* BDNF) to the culture *(Onorati, M. et al., 2016; Dell’Anno, M. et al., 2018)*.

#### 4.2 Split of cells

Cells are split as described in *(Dell’Anno, M. et al., 2018)*.

### 5. h-SC-NES cells transduction with GFP carrying lentiviral vector

#### 5.1 Preparation of the solution for cell transduction

Cell transduction medium is composed by a specific volume of NES medium and a precise volume of lentiviral stock preparation according to different parameters:

- the desired MOI (multiplicity of infection = ratio of the numbers of viral particles to the numbers of the host cells in a given infection medium)
- the number of cells we plate
- the concentration of lentiviral stock preparation
- the surface area of the used culture vessel

NOTE: We used MOI 3, based on our previous lab experience, but the MOI could vary depending on the used cell line and lentiviral preparation. If the desired MOI is = 3 and the number of plated cells is 0.5*10^5^/cm^2^ in 1W-MW24 (2cm^2^), assuming that the PFU/TU (plaque forming unit/ transducing unit) of lentiviral preparation is 25*10^6^ PFU/TU in 1ml, the calculations is as follows:

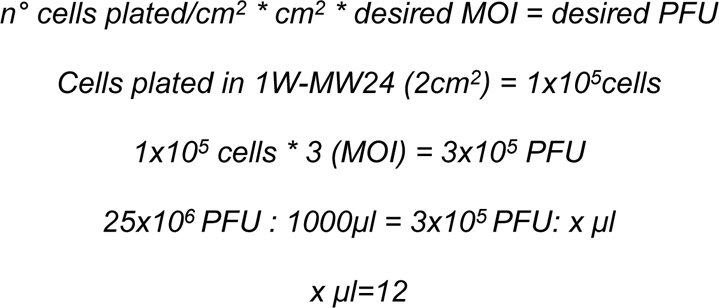

To transduce the cells with cell transduction medium-MOI3 in 1W-MW24, add 12 µl of LV preparation to NES medium (238 µl), for a total volume of 250 µl.

The medium is usually prepared fresh on the transduction day.

#### 5.2 h-SC-NES transduction protocol

1. h-SC-NES cells at a low passage are plated in POLFN-coated 24-multi well (or any other culturing support) at a density of 0.5×10^5^/cm^2^ in NES medium.
2. The next day conditioned NES medium is collected from the wells and stored. Depending on the chosen culturing support, the lowest volume of transduction medium necessary to cover the seeding surface is added to the cells (*e.g.* 150 µl /well of 24-multiwell plate). This allows to uniformly cover the well surface.
3. h-SC-NES cells are then incubated for 6h at 37°C. After that, the previously collected conditioned medium is added (200 µl for a 24-multi well) and cells are incubated O/N at 37°C.
4. The next day, h-SC-NES cells are washed once in DPBS and a total medium change (NES medium) is performed.
5. The following days cells are checked at the fluorescence microscope to observe GFP expression.
6. h-SC-NES cells could be expanded for cell banking.

### 6. Cells transplantation into SC slices and co-culturing

#### 6.1 Preparation of glass microneedles

1. Use a puller to obtain fine needles from borosilicate glass capillaries. Set the puller as follows: HEAT 990, PULL 350. NOTE: From one capillary, it is possible to obtain two fine needles.

#### 6.2 Cell preparation for transplantation

1. Split cells as described in methods section 4.2. NOTE: If your cells do not express a fluorescent reporter, label them with a cell tracking dye to monitor them at a fluorescence microscope after transplantation and during long-term culturing. Follow the chosen company protocol for the labeling step.
2. Count cells after the split and centrifuge. Suspend the obtained pellet with fresh medium + Y-27632 (10 µM) to have the desired concentration of cells per µl (usually a range between 30,000-50,000 cells µl^-1^).
3. Transfer the cell suspension to a 500 µl or 1.5 ml tube and place it on ice. Cells are ready for transplantation.

#### 6.3 Cell transplantation into organotypic slices

Human cell transplants into the mouse SC organotypic slices are performed using an air microinjector and glass microneedle.

1. Load a glass microneedle with 4 µl of cell suspension using a micropipette. NOTE: Avoid air bubble formation in the needle, since it could hamper the microinjection process. If bubbles are formed, remove them with the micropipette.
2. Place the needle in the assigned support of the microinjector and break the needle tip using the straight tweezers.
3. Before transplanting into the slices, set microinjection parameters on a calibrated glass slide by making a drop of mineral oil and microinjecting the cell suspension into the drop. The diameter of the obtained sphere of cell suspension correlates with a specific microinjection volume. Change the microinjection parameters (time, pressure) in order to obtain the desired microinjection volume.
4. Once set the right volume, microinject the cell suspension rapidly into the slices. Check at the fluorescence stereomicroscope the presence of the cells in the slices to verify that the microinjection was performed correctly. NOTE: Cell suspension could sometimes obstruct the needle: in this case, try to remove the block of cell suspension by modifying the injection parameters or load a new needle with fresh cell suspension.
5. After the transplantation, place the slices in a normal incubator at 37°C and 5% CO2 until the desired time point.

### 7. Immunofluorescence staining

#### 7.1 Day 1

1. Remove the medium at the bottom of the insert membrane and wash three times the slices with pre-warmed DPBS.
2. Fix the slices with pre-warmed 4% formaldehyde (FA): remove DPBS and add 1.5 ml of 4% FA at the bottom of the insert-membrane with the slices. After 15 min incubation at RT, add 1 ml more of 4% FA on the superior surface of the insert-membrane and incubate for 15 min at RT. Total fixation time: 30 min at RT.
3. Remove 4% FA and wash the slices three times for 10 min with DPBS.
4. Cut circumferentially the membrane of the insert with a surgical knife and separate the membrane with the slices from the plastic component of the insert. NOTE: After this step, the membrane with the slices is floating in DPBS in the dish.
5. Permeabilize with 1 ml/membrane of a solution with 0.7% Triton in DPBS for 10 min at RT.
6. Remove the permeabilization solution and incubate the samples for 4h at 4°C with 1 ml/membrane of blocking solution composed of 0.5% Triton, 10% FBS in DPBS.
7. Remove the blocking solution and add to the slices the primary antibodies at their working dilution [*e.g.*, mouse anti-Neurofilament (NFL) antibody, 1:500; rabbit anti-NFL, 1:500; rabbit anti-RBFOX3 (NeuN) antibody, 1:400; rabbit anti-active Caspase 3 (aCasp3), 1:400; mouse anti-human Nuclei, (Hu-Nu) 1:400; rabbit anti-Hu-Nu 1:400; mouse anti-GFP 1:400] within 1 ml of antibody solution composed of 0.5% Triton, 1% FBS in DPBS. Incubate overnight at 4°C.

#### 7.2 Day 2

1. Wash three times with 1-2ml of DPBS for 10 min.
2. Incubate the membrane with secondary antibody [*e.g.*, Goat anti-Mouse IgG (H+L) secondary antibody, Alexa Fluor 488, 1:500; Goat anti-Rabbit IgG (H+L) secondary antibody, Alexa Fluor 568, 1:500; Goat anti-Mouse IgG (H+L) secondary antibody, Alexa Fluor 647, 1:500; Goat anti-Rabbit IgG (H+L) secondary antibody, Alexa Fluor 647, 1:500] and HOECHST/DAPI for nuclei diluted in 1 ml/membrane of antibody solution (Triton 0.5% + FBS 1% in DPBS) for 3 h at RT. NOTE: Carefully protect the samples from the light to avoid secondary antibody bleaching during incubation and in the following steps.
3. Remove antibody solution and wash three times with DPBS (1-2 ml) for 10 min.
4. Replace DPBS with fresh one and store at 4°C in light-protected conditions; alternatively, mount the membranes on glass slides with mounting solution and leave them to dry overnight, protected from the light.
5. Store the samples at 4°C in the dark or perform imaging analysis.

### 8. Live/Dead assay

1. Prepare the working solution by aliquoting 700 µl per dish of fresh OM and add at the correct working dilution the Sytox (*e.g.*, Component B, 1:2000) and the Calcein AM (*e.g.*, Component A,, 1:2000). NOTE: Reagents are light sensitive, thus protect the working solution from light.
2. Evaluate the medium volume at the bottom of the membrane and add the Sytox and the Calcein AM at the same working dilution described at point 1.
3. Add 2 drops on the top of each slice of the working solution prepared at point 1. NOTE: Protect the dish from the light by placing them in dark conditions.
4. Incubate the slices for 30 min at RT.
5. After the incubation, cut out circumferentially the membrane from the insert with a surgical knife: after that, the membrane with the slices is floating in the working solution.
6. Mount upside down the membranes w/o the mounting solution on a cover slip and add 100 µl PBS on the top of the membranes to keep them hydrated.
7. Acquire live images at the confocal microscope as fast as possible. NOTE: Add 2 drops of PBS on the top of the membranes every 30 minutes during image acquisition to prevent the membranes from drying out.

### 9. Imaging

#### 9.1 Confocal imaging of immunostained samples

1. For qualitative analysis, acquire images at a confocal microscope using the following acquisition parameters: set the large image option (choose: 4×4), use 10X objective, no stacks, and a resolution of 3634×3634 pixels.
2. For quantitative analysis (aCasp3, Calcein and Sytox for slices and aCasp3 for cells), acquire images at confocal microscope using the following acquisition parameters: 20X objective, resolution of 1024×1024 pixels with a Z-spacing of 3 μm.

#### 9.2 Live imaging of transplanted slices at the stereomicroscope

1. Capture images at the stereomicroscope in bright-field and epifluorescence.
2. Using *bright-field* setting, acquire pictures of the slices: we used objective 1X with zoom 3X. NOTE: Modify the light depending on the used microscope and use the optical fibers if it is necessary.
3. Using *fluorescence* setting, acquire pictures of the transplanted cells with the same objective and zoom used for the slices. Parameters used for the acquisition: gain 1, exposure between 200-500 ms, offset -10.

#### 9.3 Live imaging after LIVE/DEAD assay at confocal microscope

1. Acquire images at a confocal microscope using the following acquisition parameters: 20X objective, resolution 1024×1024 pixels with a Z-spacing of 3 μm.

### 10. Picture analysis by ImageJ

#### 10.1 NFL, RBFOX3 and DAPI area analysis

1. Open ImageJ software (https://imagej.net/software/imagej/).
2. Open the file image (Click *File > open > select file > open*)).
3. In the pop-up window, select:
  - *Stack viewing > Hyperstack*
  - *Color mode > Default (with autoscale)* Select *ok* to open the image.
4. Select *Image > Color > Split channels*.
5. Choose the desired channels to analyze, *e.g*.: green channel for NFL (axonal marker), red channel for RBFOX3 (neuronal marker) and blue channel for DAPI.
6. For NFL analysis proceed with the following steps:
  - *Image > Adjust > Threshold > Select the parameters* (dark or white background, algorithm *e.g.*: default) and move the cursor on the value bar (under/over) to cover all the neurite area *> Set > Apply*.
  - Select from the tool bar the *tracing tool Wand* and use it to automatically select the area covered by NFL.
  - *Analyze > Measure > Area value in µm^2^*.
7. For DAPI and RBFOX3 analysis proceed with the following steps:
  - *Image > Adjust > Threshold > Select the parameters* (dark or white background, algorithm *e.g.*: default) and move the cursor on the value bar (under/over) to cover all the RBFOX3 or DAPI area *> Set > Apply*.
  - *Process > FFT > Bandpass Filter*.
  - Use the Threshold value bar to adjust the area covered by RBFOX3 or DAPI.
  - Select from the tool bar the *tracing tool Wand* and use it to automatically select the area covered by RBFOX3 or DAPI.
  - *Analyze > Measure > Area value in µm^2^*.

#### 10.2 Analysis of apoptosis by ImageJ

1. Open ImageJ software (https://imagej.net/software/imagej/).
2. Open the Z-stack file image (Click *File > open > select file > open*).
3. In the pop-up window, select:
  - *Stack viewing* > *Hyperstack*
  - *Color mode > Default (with autoscale)* Select *ok* to open the image.
4. Select *Image > Color > Split channels*.
5. Choose the desired channels, *e.g*.: red channel for aCasp3 (apoptosis marker to analyze) and blue or cyan for DAPI or Hu-Nu for nuclei. Then, overlay the channels selecting *Image > Color > Merge channels > select create composite > ok*.
6. Drag the Z-bar at the bottom of the image to browse through the Z-stack of the image and identify stacks in the central region of the slices with aCasp3 positivity.
7. Select *Plugins > Analyze > Cell counter*.
8. In the opened pop-up window select *Initialize* to prepare the image for the count; then select a *counter type e.g. Type 1* and rename it as the object to count (*e.g.* aCasp3^+^ cells). Rename other counters type as described above, if it is necessary to count another object (*e.g.* DAPI or Hu-Nu for the total number of cells).
9. Select in the pop-up window the counter type corresponding to the object to count, then select the *Point tool* in the tools bar and begin to count the number of apoptotic cells, positive to aCasp3.
10. Click another counter type in the cell counter window and begin to count the total number of cells (DAPI, for slices; Human Nuclei, for transplanted cells).

#### 10.3 Analysis of LIVE/DEAD assay by ImageJ

1. Open ImageJ software (https://imagej.net/software/imagej/).
2. Open the Z-stack file image (Click File > open > select file > open).
3. In the pop-up window, select:
  - *Stack viewing* > *Hyperstack*
  - *Color mode > Default (with autoscale)* Select *ok* to open the image.
4. Select *Image > Color > Split channels*.
5. Choose the desired channels, *e.g*.: green channel for Calcein (vitality marker to analyze) and cyan channel for Sytox (dead marker). Then, overlay the channels selecting *Image > Color > Merge channels > select create composite > ok*.
6. Drag the Z-bar at the bottom of the image to browse through the Z-stack of the image and identify stacks in the central region of the slices with Calcein and Sytox positivity.
7. Select *Plugins > Analyze > Cell counter*.
8. In the opened pop-up window select *Initialize* to prepare the image for the count; then select a *counters type e.g. Type 1* and rename it as the object to count (*e.g.* Calcein^+^ cells). Rename other counters type as described above, if it is necessary to count another object (*e.g.* Sytox^+^ cells).
9. Select in the pop-up window the counter type corresponding to the object to count, then select the *Point tool* in the tools bar and begin to count the number of Calcein^+^ cells
10. Click another counter type in the cell counter window and begin to count the Sytox^+^ cells.

### 10. Graphs generation by Prism (GraphPad)

All statistical analysis and graphs were obtained with Prism-Graph Pad 9 software.

## REPRESENTATIVE RESULTS

The described method allows the establishment of SC organotypic slices from mice at stage P3 and their maintenance in culture for a prolonged time in healthy conditions. Moreover, we show a protocol for transplanting cells into the slices and for co-culturing them for up to 30 days. (**Figure S1**). Firstly, we show the optimization of the culture conditions and a protocol suitable for prolonged culturing of the SC slices with transplanted cells (**Figure 1A**). Slices are generated and maintained from DEV 0 until DEV 2 in OM, which was originally proposed as an optimal medium for the maintenance of SC slices (*Vyas, A. et al., 2010*). However, due to the presence of serum proteins, this medium could be sub-optimal to sustain the neuronal differentiation and maturation of the transplanted neural precursor cells. Indeed, at DEV3, we tested the switch from OM to GM, a formulation containing Neurobasal plus B27, which supports neural survival, and without serum which inhibits the correct neuronal differentiation, promoting instead a glial fate (*Brewer, G.J. et al., 1993; De Vries, G., and Boullerne A., et al., 2010*). **Figure 1B** shows the results achieved by switching the medium at DEV3 from OM to GM, compared to the SC slices not receiving the switch (control slices were cultured in OM). We used the distribution of NFL signal inside the slices as a marker for neuronal integrity (**Figure 1B-C**). Slices at DEV 7 were healthy in both culturing conditions, showing the diffuse distribution of neurofilament (NFL, in green) inside them. At DEV 10, slices cultured in GM seemed to be healthier with respect to the control slices cultured in OM, as documented by NFL staining distribution. We also estimated NFL^+^ area (% NFL^+^ Area/DAPI^+^ Area) of the slices shown in the representative pictures of **Figure 1B**. The estimated NFL^+^ area is represented in the histograms in **Figure 1C**, confirming that NFL signal is diffusely distributed in the slices at DEV 7 in both conditions. However, at DEV 10 the estimated area covered by NFL staining decreases for OM culturing condition. These data suggest that switching to the GM at DEV3 is well-tolerated for prolonged culturing of SC slices (DEV 10). As a next step, we tested GM at more prolonged time points: DEV 30, DEV 60, and DEV 90. As shown in **Figure 2 A-B**, slices were maintained healthy in culture until DEV 90. NFL staining was found widely present in the slices at each time point, with diffuse sprouting around the slices of neurites departing from the central region. Indeed, we estimated the NFL^+^ area of the slices shown in **Figure 2A** and it increased over time as shown in the histograms of **Figure 2B**. We also observed positivity to the neuronal marker RBFOX3, providing another line of evidence of the neuronal differentiation of the slices. At each time point, we also checked the apoptosis rate by evaluating in different slices the number of cells positive to active-Caspase3 (aCasp3) (**Figure 3A-B**). The analysis was performed as described in section 9.2 of the methods. Apoptotic rate (% aCasp3^+^ cells/Total number of DAPI^+^ cells) was found to be very low at each time point (0.85±0.52, 0.71±0.27, 0.66±0.45 for DEV 30, 60, and 90, respectively) with no statistically significant differences between the three considered time points (*Kruskal-Wallis Test, multiple comparisons, p-value>0.05*, **Figure 3B**). These data suggest that the apoptotic rate associated with aCasp3 remains stable during time and, together with the wide distribution of NFL into the slices (**Figure 3A**), confirms the survival of the slices at each time point.

**Figure 1.**
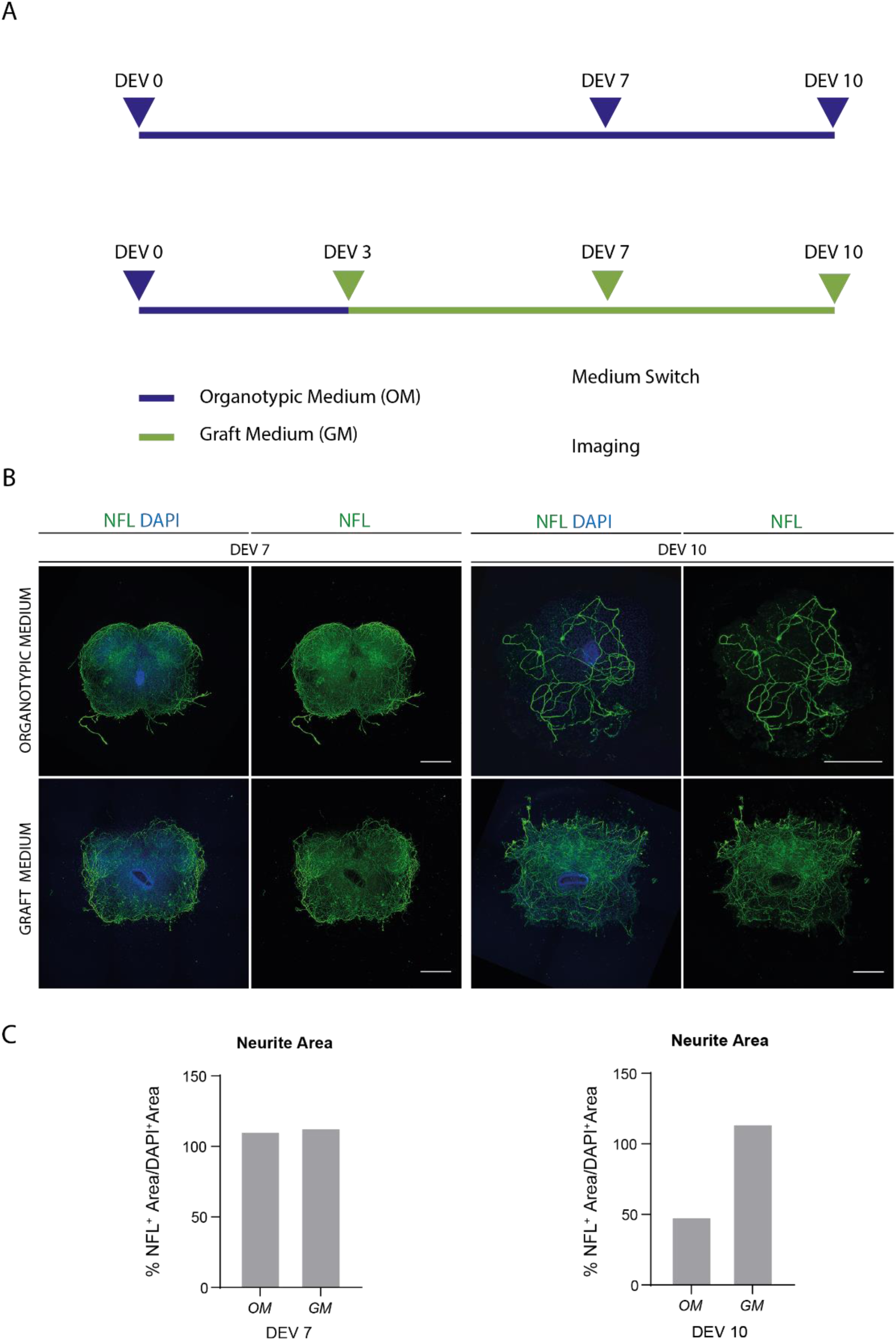
Optimization of long term culturing conditions. **(A)** Representative scheme of protocol for testing organotypic medium (OM) and graft medium (GM). OM was maintained until DEV 7-10 for control group. The medium was switched to the GM at DEV 3 for the treated slices, then they were fixed at DEV 7-10 for comparison to controls. **(B)** Representative pictures comparing mouse SC organotypic slices at DEV 7 and 10 cultured in different conditions. Slices were stained for the cytoskeletal marker neurofilament (NFL, green). The wide distribution of NFL staining in slices cultured with GM suggests an overall survival and differentiation. Nuclei were counterstained with DAPI. Scale Bar: 500 µm. **(C)** Representative histograms of the estimate of area covered by NFL in the slices showed in Figure 1B. At DEV 10, NFL surface area decreases in the OM culturing condition.

**Figure 2.**
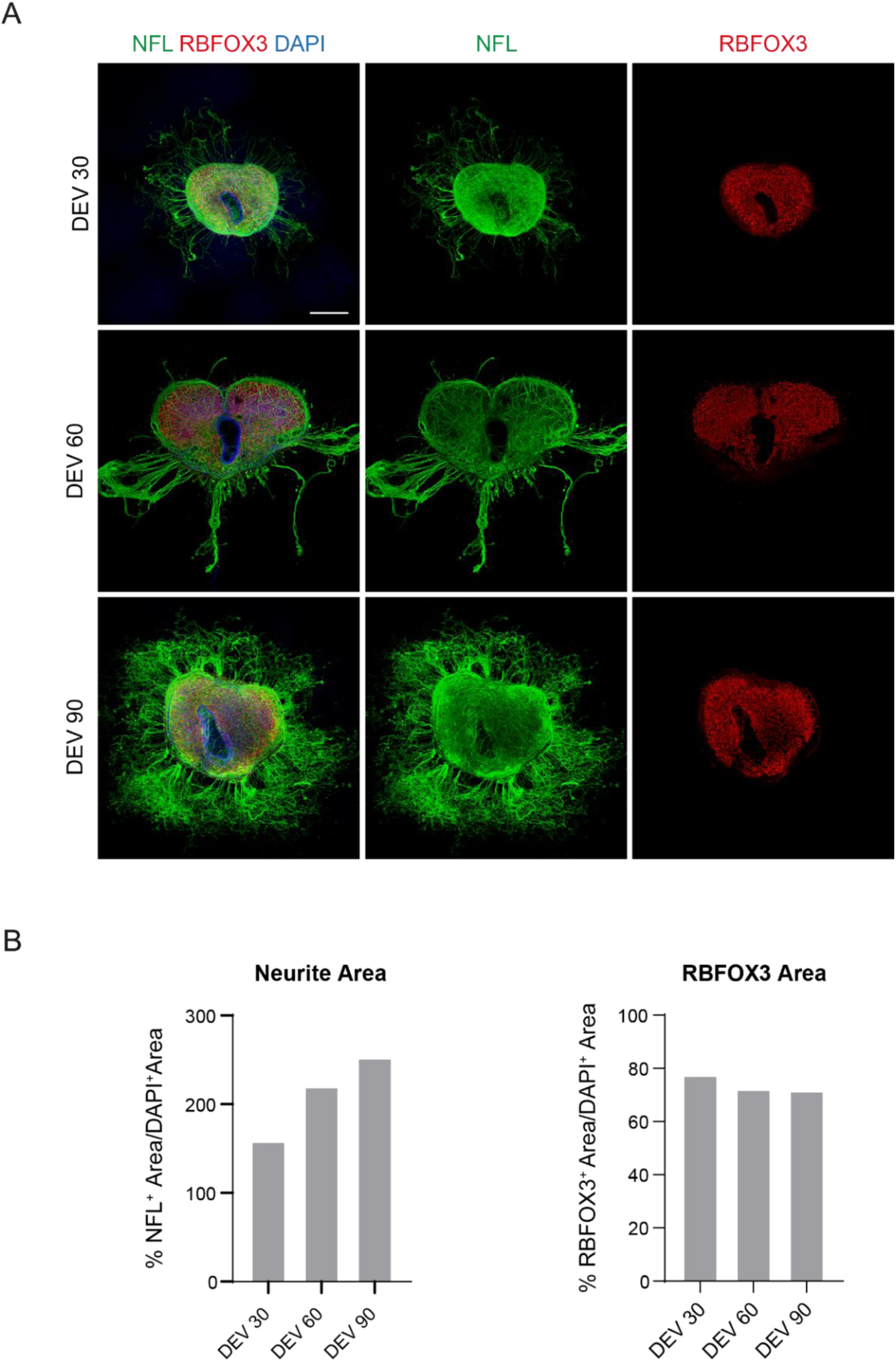
Long term cultured mouse SC organotypic slices. **(A)** Slices were maintained in culture until day *ex vivo* (DEV) 90. Immunofluorescence assay revealed wide distribution of the cytoskeletal marker neurofilament (NFL, green) and of the nuclear neuronal marker RBFOX3 (red), attesting their healthy condition and neuronal identity after long term culturing. Of note, NFL^+^ axons sprout out diffusely around the slices over time. Nuclei were counterstained with DAPI. Scale Bar: 500 µm. **(B)** Representative histograms of the estimate of NFL^+^ and RBFOX3^+^ areas of the slices showed in Figure 2A. NFL^+^ neurite area increases over time.

**Figure 3.**
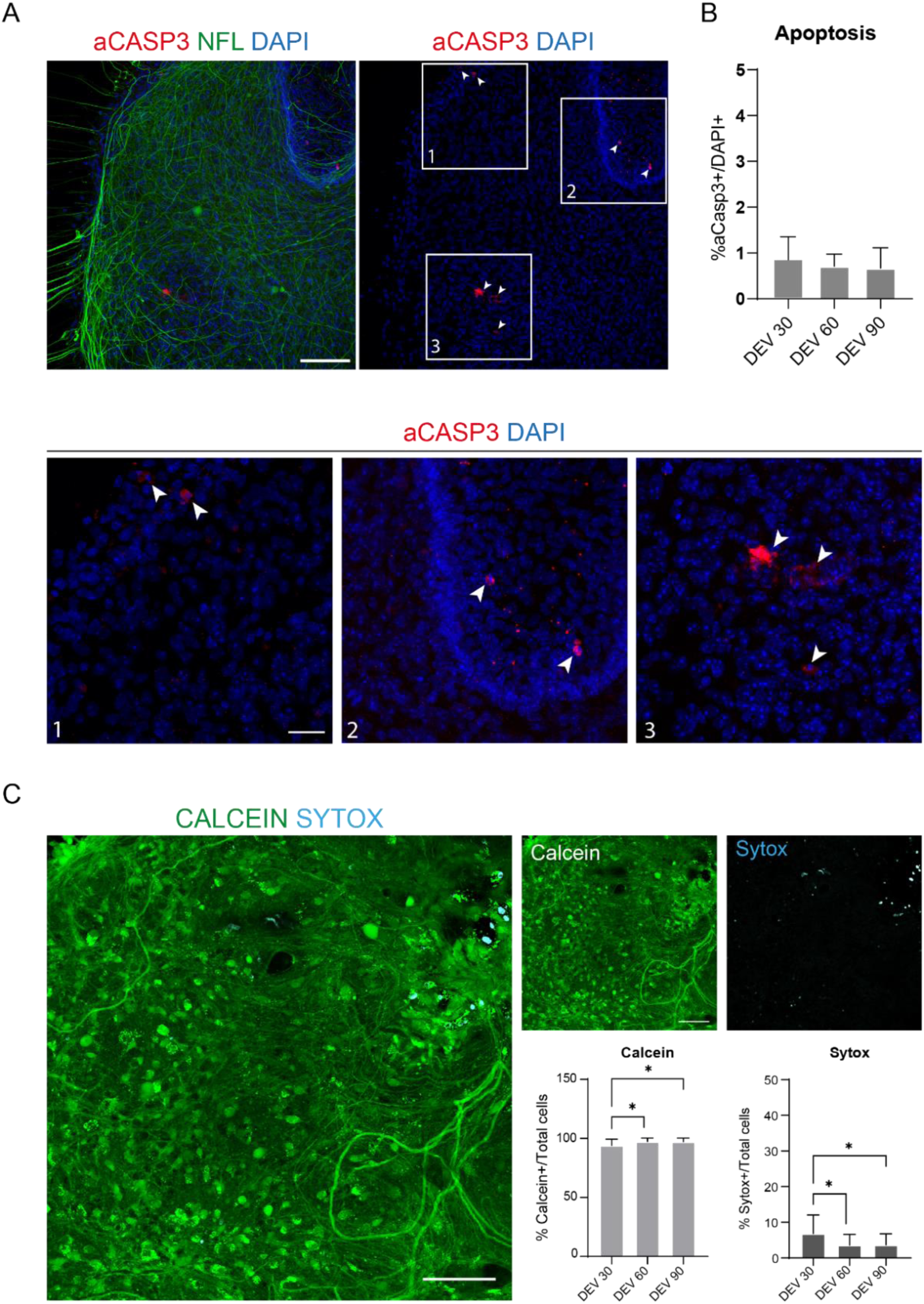
Evaluation of cell viability in the SC slices over time. (**A**) Representative pictures of organotypic slices at DEV 60 stained for aCasp3 (red) and NFL (green). Scale Bar: 100 µm. In the pictures, it is possible to observe the NFL diffuse pattern. Rare cells are positive for the apoptotic marker aCasp3 (Insets: 1-2-3). (**B**) Analysis of the apoptosis rate in slices at different time points. Mean *± SD*, N (replicates) = 6 slices, n (total cells) > 1000 for each slice, *Kruskal-Wallis test, multiple comparison, p-value > 0.05*. The apoptotic rate is stable over time. In the insets 1-2-3 of Figure 3A it is possible to observe more in detail cells positive to aCasp3 (red staining, white arrows). Small red dot label cell debris and piknotic nuclei. Scale Bar: 50 µm. (**C**) Representative pictures of live/dead assay performed on SC slices at DEV 90: metabolically active cells are labelled in green with Calcein, while dead and damaged cells are labelled in light blue (cyan) with Sytox. The two histograms show the % of cells positive to Calcein (on the left) and Sytox (on the right) on the total number of cells. For both mean *± SD*, N (replicates) = 6 slices, n (total cells) > 1000 for each slice, *Kruskal-Wallis test, multiple comparison,* DEV 30 vs DEV 60 p-value=*0,018*; DEV 30 vs DEV 90 p-value=*0,027*; DEV 60 vs DEV 90 p-value *>0,99*.

In support of previous data, we also performed a live/dead assay to evaluate the viability of the slices at the three different time points. We used Calcein (green staining) to label the viable and metabolically active cells and Sytox (cyan staining) to assess cell death. As shown in the histograms in **Figure 3C**, the percentage of metabolically active cells increases slightly from DEV 30 to DEV 90 (93.17±5.21, 96.43±3.02, 96.33±3.10 for DEV 30, 60, and

90, respectively), stabilizing between the last two time points (*Kruskal-Wallis test, multiple comparisons, DEV 30 vs DEV 60 p-value=0.018; DEV 30 vs DEV 90 p-value=0,027; DEV 60 vs DEV 90 p-value = 0,99)*. We found low levels of cell death that decreased over time (6.83±5.21, 3.57±3.02, 3.66±3.10 for DEV 30, 60, and 90, respectively) and a statistically significant difference was found between DEV 30 and later time points, DEV 60 and DEV 90 (*Kruskal-Wallis test, multiple comparisons, DEV 30 vs DEV 60 p-value=* 0,018*; DEV 30 vs DEV 90 p-value=* 0,027*; DEV 60 vs DEV 90 p-value = 0,99*). These data, in association with the apoptosis rate, confirm slice survival over time and support the effectiveness of the long-term culturing protocol performed.

Once established the ability of prolonged culturing of the SC slices, we challenged the system by transplantation of h-SC-NES cells at the first stages of neuronal differentiation. We tested the h-SC-NES cells because they have shown promising results for SCI treatment (*Dell’Anno, M. et al., 2018*). The transplantation procedure of h-SC-NES cells into the mouse SC slices is described in section 5 of the methods. The SC slices and transplanted h-SC-NES cells were maintained until DPT 30. Cells were grafted at DIV 10 of differentiation (neural precursor stage) into DEV 4 organotypic slices, as shown in the protocol scheme of **Figure 4A**. Transplanted cells were monitored for the expression of GFP in culture for up to 30 days. **Figure 4B** shows representative live images, at different DPT, of a SC slice with transplanted GFP^+^ cells. The stable expression of GFP over time (**Figure 4B** and **Figure 5A**) suggests that cells survived to prolonged grafting into SC tissue in the previously optimized culture conditions. We also checked the apoptotic rate of transplanted cells as described in section 9.1 of the method. Apoptotic rate (% aCasp3^+^ cells/Total number of Hu-Nu^+^ cells) was found to be very low (0.44±0.34 %) after 30 DPT (**Figure 5B**). Moreover, the apoptotic rate at DPT 30 was found to be in line with that one found for the same type of cells at DPT 7, reported in *De Vincentiis, S. et al., 2023*, documenting that the cultures stabilize with time.

**Figure 4.**
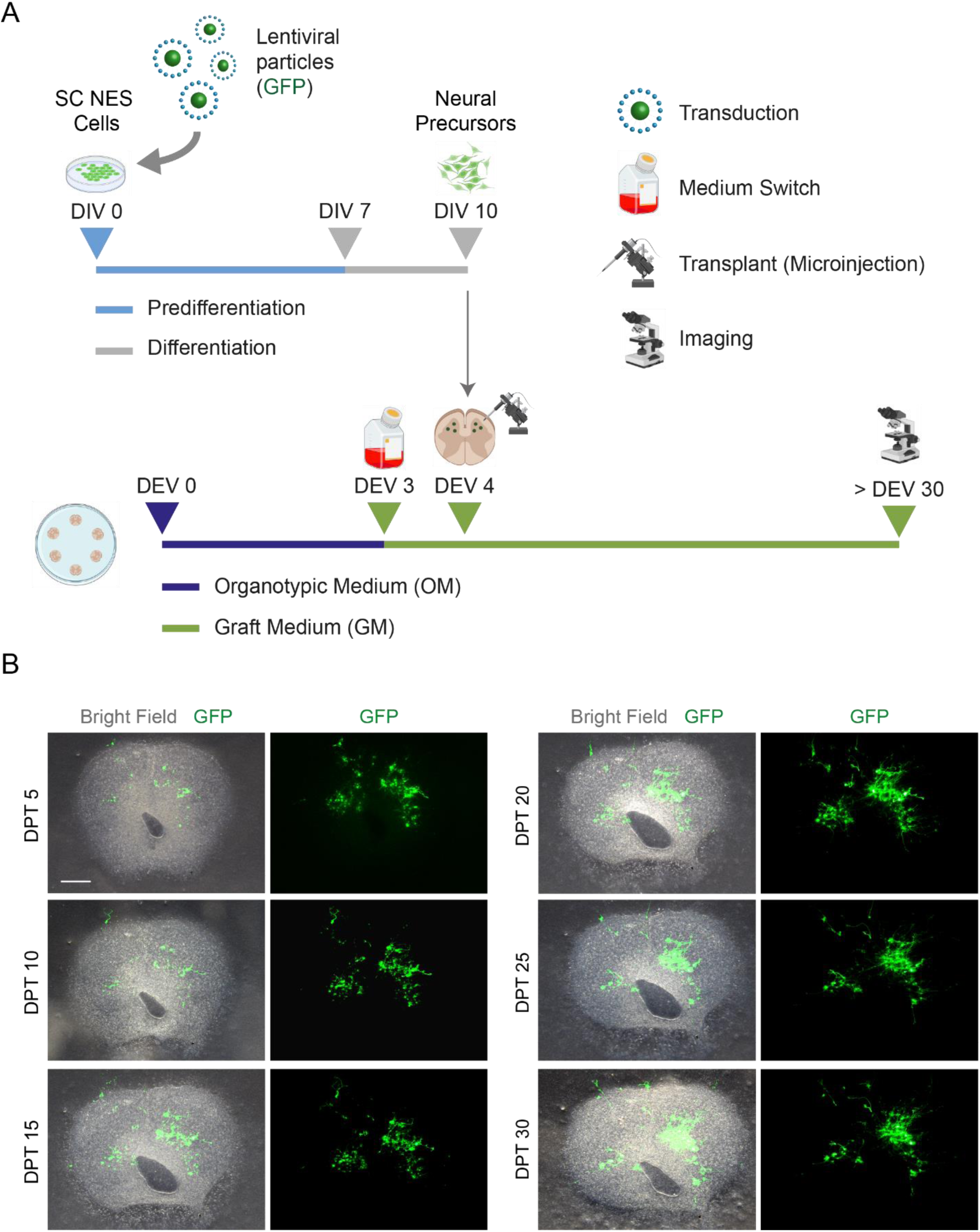
Human SC-NES cell transplantation into mouse organotypic slices. **(A)** Representative scheme of transplantation protocol. Cells were transplanted as neural precursors at DIV 10 of differentiation into DEV 4 organotypic slices. **(B)** Representative images of mouse organotypic slices transplanted with GFP expressing h-SC-NES cells over time until day post transplantation (DPT) 30. Cells were transduced with a lentiviral vector carrying the GFP gene. GFP expression over time confirmed their viability and adaptation to the slice environment. Scale Bar: 500 µm.

**Figure 5.**
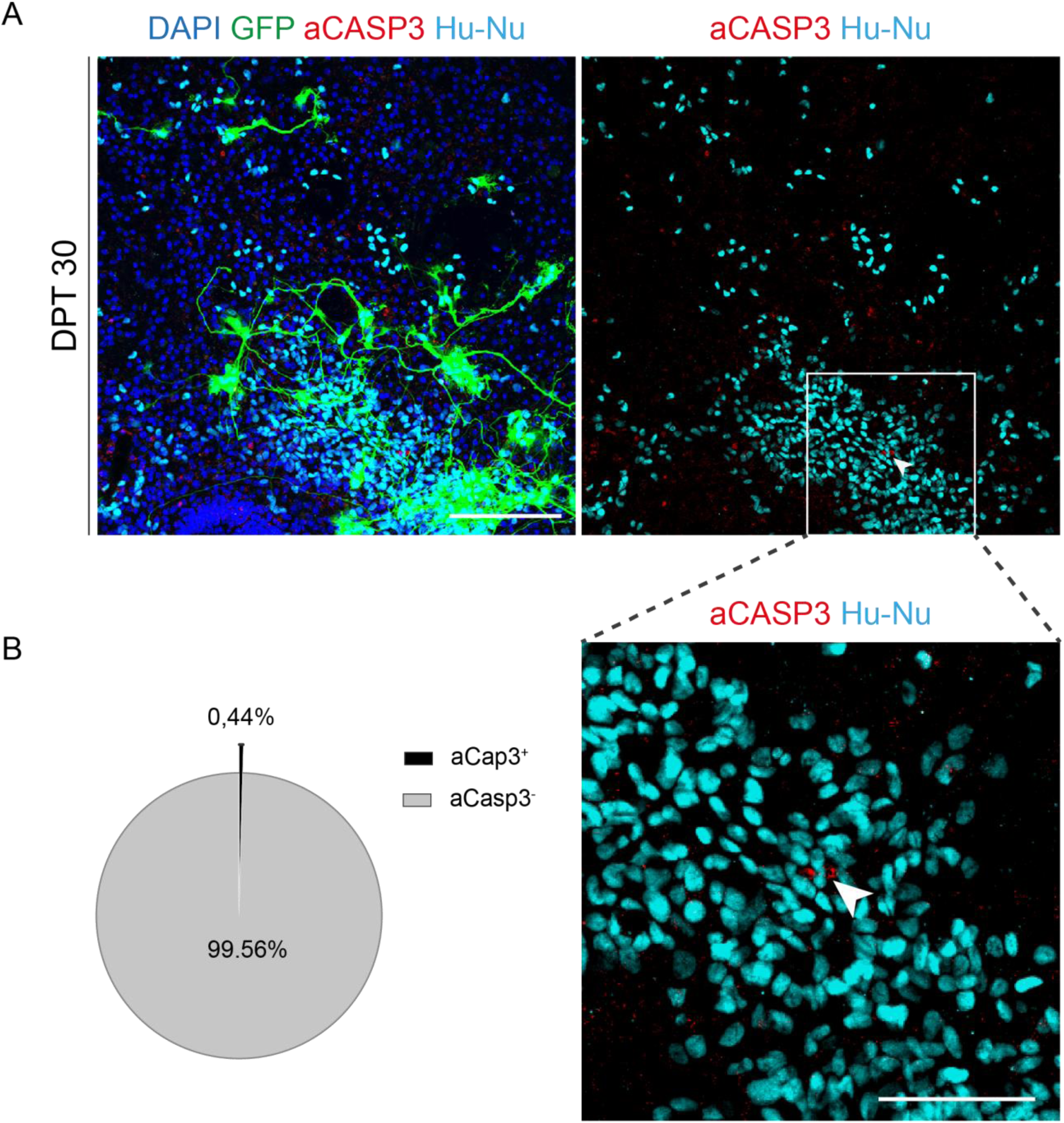
Apoptosis rate evaluation of transplanted h-SC-NES cells after 30 days from transplant. **(A)** Representative picture of a mouse organotypic slice transplanted with GFP expressing h-SC-NES cells. Cells were transduced with a lentiviral vector carrying GFP gene for monitoring them into the slices after transplantation. GFP expression over time confirmed their viability and adaptation to the slice environment. The time point shown is DPT 30; cells were stained for human nuclei (cyan) and aCasp3 (red). Scale Bar: 150 µm. **(B)** On the left, representative pie chart of the apoptosis analysis of cells transplanted in slices at DPT 30 (N (replicates) = 5 slices, n (cells) = 5000) and on the right, an inset of Hu-Nu positive cells and a detail of a cell positive to aCasp3 (white arrow). Scale bar 75 µm. Small red dots label cell debris and piknotic nuclei.

## DISCUSSION

There is still no effective treatment for patients with SCI. Different approaches have been tested and one of the most promising is based on a regenerative strategy, i.e. cell replacement. Currently, the advancements in the regenerative medicine field ask for novel platforms to test the efficacy and safety of cell transplants, alone or in combination with other approaches. Their preclinical validation is essential to pursue further clinical studies. SC organotypic cultures are a useful platform for studying different aspects of neurodegeneration, neural regeneration and neurodevelopment and for investigating the effectiveness of novel therapeutic approaches (*Pandamooz, S. et al., 2016*). In particular, specific features of the organotypic cultures such as the maintenance of original histoarchitecture, and of cell and microenvironment composition are advantageous to unravel transplantation dynamics, such as cell engraftment, integration, differentiation, and maturation. Consistently with published protocols, SC organotypic slices can be maintained in culture for approximately 2-3 weeks in healthy conditions and this limits their use for the long-term investigations and functional screening required for testing schemes of cell therapy. Exploring important processes like differentiation and maturation towards the correct fate of transplanted cells inside SC tissue requires indeed long-term monitoring. These cellular processes are critical during common transplants in animal models. The availability of an *ex vivo* system that mimics many features present *in vivo* would be helpful in the pre-clinical screening phase. For this reason, in this work, we propose an optimal long-term SC organotypic culture method that allows to maintain viable SC slices for up to 90 days, triplicating their usual culture time frame. Moreover, we show stable h-SC-NES cell engraftment inside SC slices and the maintenance of the transplant culture for up to 30 days. We were able to monitor cell engraftment over time by observing GFP expression and to verify cell survival at DPT 30. This improved long-term *ex vivo* platform alone and in the transplant configuration will help researchers in pre-clinical screening for stem cell-based transplants for SCI. This will allow them to identify the best cell candidate for further *in vivo* studies promoting the success of the transplants. Moreover, after initial screening, SC organotypic slices could also be used in parallel to the *in vivo* studies, to confirm and corroborate long term cellular dynamics and behaviors observed in animal models or to support mechanistic studies.

Our protocol describes in detail how to generate this long-term organotypic model, but some critical steps should be also discussed. Concerning the generation of the SC organotypic cultures, there are some challenges during the surgery and the first stages of culture. A well-performed surgery procedure is essential to generate slices that maintain the original cytoarchitecture. If the SC is ruined during the isolation, slices can lose their typical anatomic structure and tissue damage can induce an excessive pro-inflammatory insult leading to unhealthy conditions and cell death. The most challenging phase during the surgery is the extraction of the SC from the backbone and the removal of meninges from the isolated SC. The success of these steps depends on the experience of the operator, thus a training period before starting with the experiments is recommended. Coronal sectioning of the SC through a chopper is also a challenging phase. The SC should be placed on the cutting deck exactly perpendicular respect to the blade. The operator should also place the blade perpendicularly to the cutting deck. These precautions are necessary to ensure the generation of reproducible slices among the same and different experiments. Another important issue is that the time for surgery is limited: the entire slice generation procedure must take around 30 minutes. If the operator spends more time on surgery and cutting, the SC tissue will suffer and this can impair the success of the culture and the next steps of the experiment. Once slices are placed on the culture membrane, it is important to feed them correctly. GDNF is necessary to sustain tissue recovery and survival. Cutting with a chopper is traumatic for the tissue and, for this reason, slices are placed soon after the cut into an ice-cold dissection medium to clean away pro-inflammatory and death-promoting molecules. Then, slices are placed on the culture membranes (cell culture inserts) with fresh medium modified with GDNF to promote a faster recovery and slice adhesion to the membrane. GDNF should be added to the medium every day for the first week in culture, because of its short half-life (*Ziv-Polat, O. et al., 2014; Mesa-Infante, V. et al., 2022*). We observed that slices need the continuative presence of GDNF during the first days in culture to promote tissue recovery and viability. In any case, GDNF presence is important for the entire culturing period, thus, it is strongly discouraged to interrupt GDNF administration at further time points. During the first week in culture, it is also important to check the slices macroscopically by eye and at the microscope. Translucent tissue and transparency of the borders are signs of proper adhesion of the slices to the membrane and of viable tissue. The necrotic tissue will appear extremely white at first macroscopic sight and the necrotic areas will appear dark grey at the microscope. After some weeks in culture, the morphology of tissue may change: cell movements and tissue adhesion to the membrane can influence this process. We observed, for example, in some slices the loss of the central lumen, filled with cells, and the loss of dorsal and ventral horn morphology. This happens mainly with smaller slices, while most of the slices will maintain an anatomic structure close to the original one. Concerning transplanting cells into the slices, the main issue is related to the break of the glass microneedle tip. If the hole for the passage of cells is too big, it can cause damage to SC tissue during the microinjection. If it is too small, cell stacking can obstruct the needle, hampering the transplantation process. The transplantation procedure should be completed within 1h, to minimize cell suffering and death.

In conclusion, the proposed protocol ensures obtaining an optimal and versatile tool for different types of investigations. Here, we apply our long-term platform to validate the transplantation of h-SC-NES cells at the first stages of differentiation inside mouse SC tissue for 30 days. These long-term organotypic cultures could also be exploited for testing neuroprotective and neuroregenerative agents or novel molecules/materials or to study neurodegenerative disorders that involve the SC. Being explants of a specific organ, organotypic cultures present features that bridge the gap between 2D cell cultures and *in vivo* models, confirming them as an invaluable tool for both basic research and pre-clinical testing.

## ACKNOWLEDGMENTS

The study was supported by the Wings for Life Foundation (WFL-IT-20/21), the European Union Next-GenerationEU - National Recovery and Resilience Plan (NRRP) – mission 4 component 2, investiment n. 1.4 – CUP N. B83C22003930001 (Tuscany Health Ecosystem – THE”, Spoke 8), and the Marina Romoli Onlus. This manuscript reflects only the authors’ views and opinions, neither the European Union nor the European Commission can be considered responsible for them.

Data and metadata are available on Zenodo **10.5281/zenodo.10433147.**

Images generated with Biorender https://www.biorender.com/.

