## Supplementary figures and images for "Long-term mouse spinal cord organotypic slice culture as a platform for validating cell transplantation in spinal cord injury"

### Supplementary Figure S1

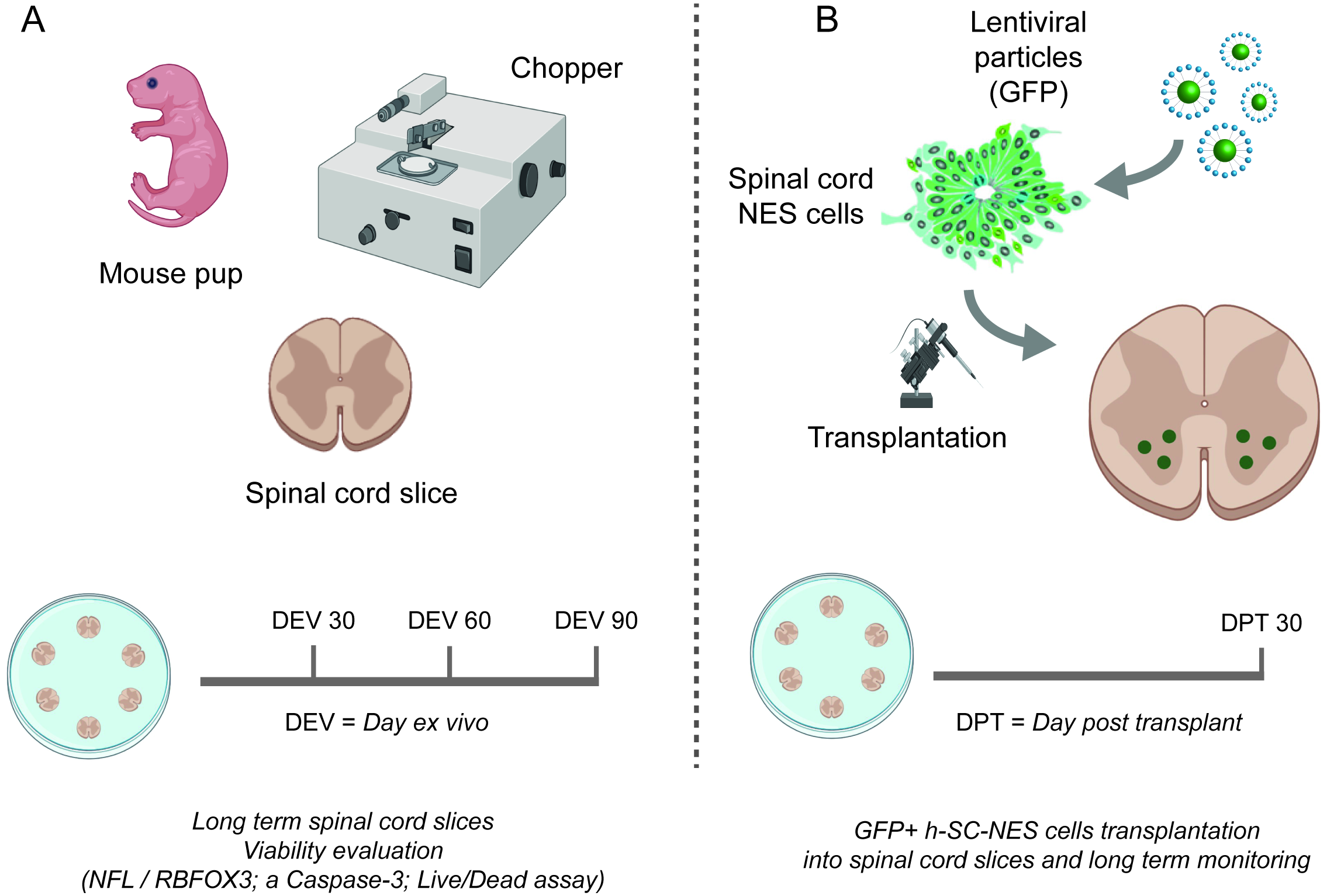
